# A central amygdala-globus pallidus circuit conveys unconditioned stimulus information and controls fear learning

**DOI:** 10.1101/2020.04.28.066753

**Authors:** Jacqueline Giovanniello, Kai Yu, Alessandro Furlan, Gregory Thomas Nachtrab, Radhashree Sharma, Xiaoke Chen, Bo Li

## Abstract

The central amygdala (CeA) is critically involved in a range of adaptive behaviors. In particular, the somatostatin-expressing (Sst^+^) neurons in the CeA are essential for classic fear conditioning. These neurons send long-range projections to several extra-amygdala targets, but the functions of these projections remain elusive. Here, we found in mice that a subset of Sst^+^ CeA neurons send projections to the globus pallidus external segment (GPe), and constitute essentially the entire GPe-projecting CeA population. Notably, chronic inhibition of GPe-projecting CeA neurons completely blocks auditory fear conditioning. These neurons are selectively excited by the unconditioned stimulus (US) during fear conditioning, and transient inactivation or activation of these neurons during US presentation impairs or promotes, respectively, fear learning. Our results suggest that a major function of Sst^+^ CeA neurons is to represent and convey US information through the CeA-GPe circuit, thereby instructing learning in fear conditioning.

## Introduction

The central amygdala (CeA) plays important roles in learning and executing adaptive behaviors. In particular, its function in the acquisition and expression of defensive behaviors has received arguably the most intensive study (Duvarci and Pare, 2014; Herry and Johansen, 2014; Janak and Tye, 2015). For example, transient pharmacological inactivation of the CeA (Goosens and Maren, 2003; Wilensky et al., 2006), or specific inactivation of the lateral division of the CeA (CeL) (Ciocchi et al., 2010), during Pavlovian fear conditioning blocks the formation of fear memories. Moreover, *in vivo* single unit recording demonstrates that fear conditioning causes increased spiking in one CeA population (the “ON” neurons) and decreased spiking in another (the “OFF” neurons) in response to cues predicting shocks. Such learning-induced changes in the responsiveness of CeA neurons to CS presentations may facilitate the expression of learned defensive responses, including conditioned freezing behavior (Ciocchi et al., 2010; Duvarci et al., 2011; Haubensak et al., 2010). These findings have led to the notion that the CeA, including the CeL, is essential for the formation of aversive memories.

The CeA is a striatal-like structure that contains medium spiny neurons mainly derived from the lateral ganglionic eminence during development (Cassell et al., 1999; Garcia-Lopez et al., 2008; Swanson and Petrovich, 1998; Waraczynski, 2016). These neurons show considerable heterogeneity (Fadok et al., 2018; Li, 2019), which is partly revealed by the different genetic or neurochemical markers that these neurons express. Two of these markers, somatostatin (Sst) (Cassell and Gray, 1989) and protein kinase C-δ (PKC-δ) (Haubensak et al., 2010), label two major populations that are largely nonoverlapping and together constitute about 90% of all neurons in the CeL (Haubensak et al., 2010; Li, 2019; Li et al., 2013).

Recent studies have shown that the excitatory synaptic transmission onto Sst-expressing (Sst^+^) CeL neurons is potentiated, whereas that onto Sst-negative (Sst^−^) CeL neurons (which are mainly PKC-δ^+^ neurons) is weakened by fear conditioning (Ahrens et al., 2018; Hartley et al., 2019; Li et al., 2013; Penzo et al., 2014; Penzo et al., 2015). Consistently, *in vivo* fiber photometry (Yu et al., 2016) or single unit recording (Fadok et al., 2017) studies demonstrate that Sst^+^ CeL neurons show increased excitatory responses to shock-predicting cues following fear conditioning, and the responses correlate with freezing behavior (Fadok et al., 2017). Moreover, inhibition of Sst^+^ CeL neurons during fear conditioning using chemogenetic (Li et al., 2013; Penzo et al., 2015), optogenetic (Li et al., 2013) or molecular (Yu et al., 2017) methods, which can abolish the fear conditioning-induced potentiation of excitatory synapses onto these neurons (Li et al., 2013; Penzo et al., 2015), impairs the formation of fear memories. These studies provide compelling evidence that Sst^+^ CeL neurons constitute an important element of the circuitry underlying fear conditioning.

In light of previous findings about the organization of CeA circuit (Duvarci and Pare, 2014; Fadok et al., 2018; Herry and Johansen, 2014; Li, 2019), Sst^+^ CeL neurons can potentially influence fear conditioning via their inhibitory interactions with other neurons locally within the CeL and the resulting disinhibition of the CeM (Ciocchi et al., 2010; Li et al., 2013), a structure that has been shown to control the expression of freezing behavior during fear conditioning through interactions with the midbrain periaqueductal gray (PAG) (Davis, 2000; Duvarci et al., 2011; Fadok et al., 2017; Krettek and Price, 1978; LeDoux et al., 1988; Tovote et al., 2016; Veening et al., 1984). Alternatively, or in addition, as Sst^+^ CeL neurons also project to many areas outside of the CeA (Ahrens et al., 2018; Fadok et al., 2018; Li, 2019; Penzo et al., 2014; Steinberg et al., 2020; Ye and Veinante, 2019; Yu et al., 2017; Zhou et al., 2018), these neurons may influence fear conditioning through their long-range projections to extra-CeA structures.

Here, we discovered that a subset of Sst^+^ CeA neurons send projections to the globus pallidus external segment (GPe), a basal ganglia structure that is best known for its role in motor control (Kita, 2007; Wallace et al., 2017) but has also been implicated in regulating emotions or affects, including fear or threat, in both humans and animals (Baumann et al., 1999; Binelli et al., 2014; Blanchard et al., 1981; Critchley et al., 2001; Hattingh et al., 2012; Hernadi et al., 1997; Ipser et al., 2013; Kertes et al., 2009; Murphy et al., 2003; Shucard et al., 2012; Sztainberg et al., 2011; Talalaenko et al., 2006). Furthermore, through *in vivo* fiber photometry and molecular and optogenetic manipulations, we revealed that this previously unknown Sst^CeA-GPe^ circuit has a critical role in representing the aversive stimulus and instructing learning during fear conditioning.

## Results

### CeA to GPe projections originate from Sst^+^ neurons

It has been reported that the CeA sends projections to the GPe (Shinonaga et al., 1992). We started to verify this result by using a retrograde tracing approach (Figure 1A). We injected a retrograde adeno-associated virus (AAVrg) encoding the Cre recombinase (AAVrg-Cre) into the GPe of *LSL-H2B-GFP* reporter mice (He et al., 2012), which express the fluorescent protein H2B-GFP (nuclear GFP) in a Cre-dependent manner. This approach led to the labeling of many neurons in the CeA (Figure 1B), confirming the existence of the CeA-GPe pathway.

**Figure 1.**
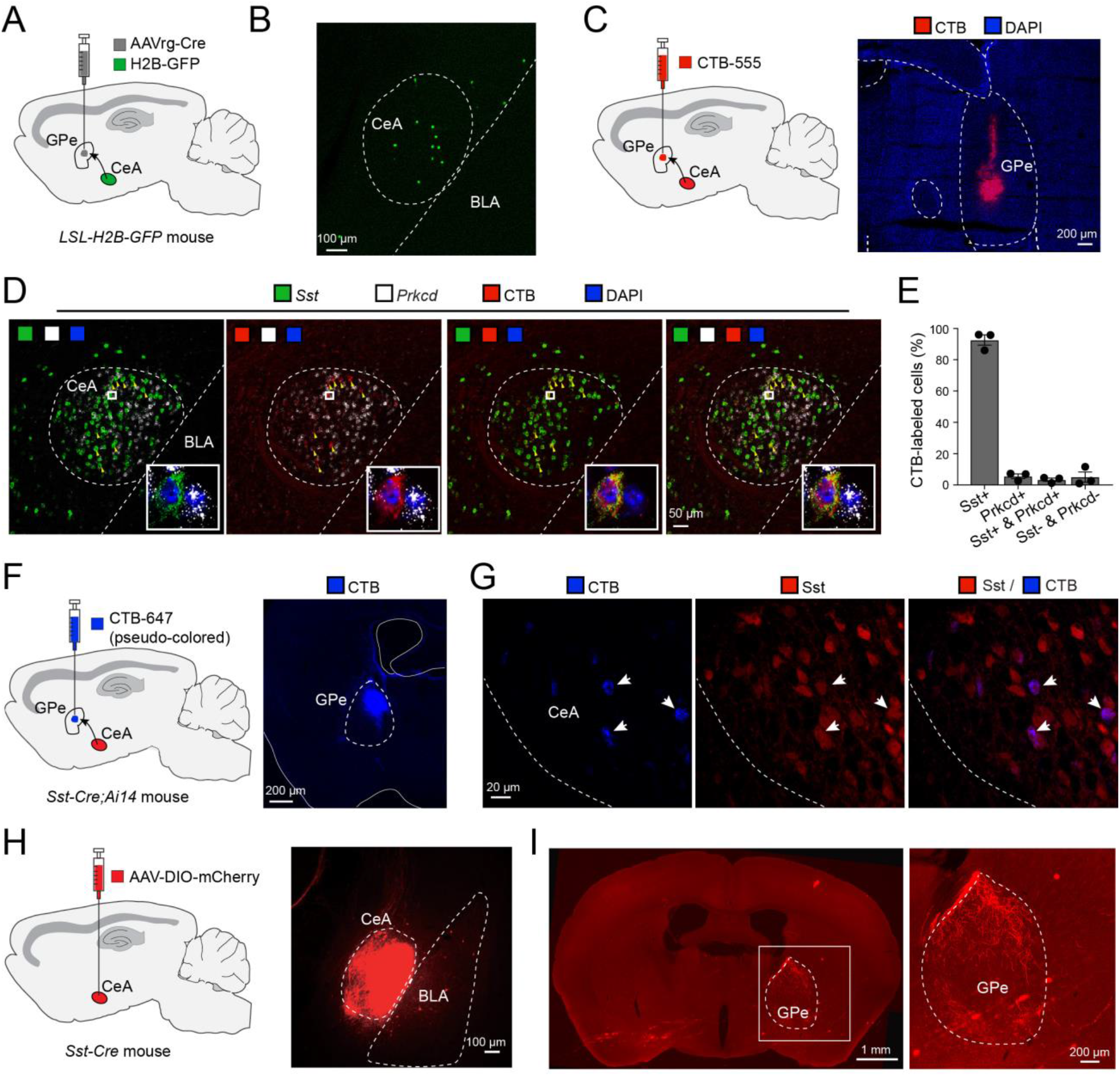
CeA to GPe projections originate from Sst^+^ neurons. (A, B) A schematic of the approach (A) and a representative image showing the retrogradely-labeled H2B^+^ cells in the CeA (B; n = 2 mice). (C) A schematic of the approach (left) and a representative image showing the target area of CTB injection in the GPe (right). (D) Confocal images of a coronal brain section containing the CeA from a representative mouse in which CTB was injected into the GPe (C), showing the distribution of GPe-projecting CeA neurons labeled with CTB, and the distribution of *Sst* and *Prkcd* expression detected with smFISH. Insets: high magnification images of the boxed areas in each of the images. (E) Quantification of the percentage distribution of different types of CeA neurons that project to the GPe (data are presented as mean ± s.e.m., n = 3 mice). (F) A schematic of the approach (left) and a representative image showing the target area of CTB injection in the GPe (right). (G) Confocal images of a coronal brain section containing the CeA from a representative *Sst-Cre;Ai14* mouse in which CTB was injected into the GPe (F), showing the distribution of GPe-projecting CeA neurons labeled with CTB, and the distribution of Sst^+^ neurons labeled with tdTomato. (H) A schematic of the approach (left) and a representative image showing the viral infection of Sst^+^ CeA neurons (red; right). (I) Left: an image of a coronal brain section containing the GPe from a representative *Sst-Cre* mouse in which Sst^+^ CeA neurons were labeled with mCherry (H). Right: a higher magnification image of the boxed area in the left, showing the distribution of axon fibers in the GPe that originate from Sst^+^ CeA neurons. This experiment was repeated in 3 mice.

To determine the main composition of CeA neurons projecting to the GPe, we injected the GPe in wild-type mice with the retrograde tracer cholera toxin subunit B (CTB) conjugated with Alexa Fluor™ 555 (CTB-555) (Figure 1C). We subsequently assessed the expression of *Sst* and *Prkcd* (which encodes PKC-δ) in the CTB-labeled GPe-projecting CeA neurons using single molecule fluorescent *in situ* hybridization (smFISH) (Figure 1D). This approach revealed that the vast majority of GPe-projecting CeA neurons expresses *Sst* (93±3%; mean±s.e.m.), whereas only a small portion of these neurons expresses either *Prkcd* (6±1%) alone, both *Sst* and *Prkcd* (3±1%), or neither of these molecules (5±3%) (Figure 1E). Similarly, retrograde tracing with CTB in *Sst-IRES-Cre;Ai14* mice, in which Sst^+^ cells are labeled with the fluorescent protein tdTomato (Madisen et al., 2010), showed that almost all the GPe-projecting CeA neurons are Sst^+^ (92±2%; n = 4 mice) (Figure 1F, G).

In a complimentary experiment, we visualized the CeA-GPe pathway using an anterograde tracing approach. An adeno-associated virus (AAV) expressing the fluorescent protein mCherry in a Cre-dependent manner was injected into the CeA of *Sst-IRES-Cre* mice to label Sst^+^ CeA neurons (Figure 1H). Four to five weeks later, we examined the brain sections from these mice for axon fibers originating from the infected Sst^+^ CeA neurons. Dense fibers were identified in the dorsal part of the GPe (Figure 1I). Together, these results demonstrate that projections from the CeA to the GPe originate predominantly from Sst^+^ neurons (hereafter referred to as Sst^CeA-GPe^ neurons).

Next, we examined the functional connectivity between Sst^CeA-GPe^ neurons and the GPe (Figure S1). We introduced the light-gated cation channel channelrhodopsin (ChR2) selectively into Sst^+^ CeA neurons of *Sst-IRES-Cre* mice, and used these mice to prepare acute brain slices containing the GPe, in which we recorded synaptic responses in neurons in response to light-simulation of the axons originating from Sst^CeA-GPe^ neurons (Figure S1A, B). About half of the neurons (5 out of 12) recorded in the GPe showed fast light-evoked inhibitory synaptic responses (Figure S1C), indicating that Sst^CeA-GPe^ neurons provide monosynaptic inhibition onto a subset of GPe neurons.

It is known that Sst^+^ CeA neurons send projections to many downstream structures (Ahrens et al., 2018; Fadok et al., 2017; Li, 2019; Penzo et al., 2014; Ye and Veinante, 2019; Yu et al., 2017; Zhou et al., 2018). Therefore, we examined whether Sst^CeA-GPe^ neurons send collateral projections to another major target of the CeA, the bed nucleus of the stria terminalis (BNST), because our recent study shows that BNST-projecting CeA neurons are also predominantly Sst^+^, and these neurons play a critical role in anxiety-related behaviors (Ahrens et al., 2018). To this end, we injected both the GPe and the BNST in the same mice with CTB conjugated with different fluorophores, such that GPe-projecting neurons and BNST-projecting neurons in the CeA were labeled with distinct colors (Figure S2A-C). Notably, we found almost no doubly labeled neurons in the CeA in these mice (<1%; Figure S2D), indicating that Sst^CeA-GPe^ neurons and Sst^CeA-BNST^ neurons are distinct populations.

### Sst^CeA-GPe^ neurons are necessary for fear learning

As both Sst^+^ CeA neurons (Fadok et al., 2018; Li, 2019) and the GPe (Blanchard et al., 1981; Hattingh et al., 2012; Ipser et al., 2013; Kertes et al., 2009; Murphy et al., 2003; Sztainberg et al., 2011; Talalaenko et al., 2006) have been implicated in processing negative affects including fear, we set out to examine the role of Sst^CeA-GPe^ neurons in Pavlovian fear conditioning. To determine whether Sst^CeA-GPe^ neurons are necessary for fear conditioning, we selectively blocked neurotransmitter release from these neurons with the tetanus toxin light chain (TeLC) (Murray et al., 2011). To this end, we used an intersectional viral strategy in wild-type mice, in which we bilaterally injected the GPe with the AAVrg-Cre and the CeA with an AAV expressing TeLC-GFP, or GFP (as the control), in a Cre-dependent manner (Figure 2A, B). Four weeks following viral injection, both the TeLC group and the GFP control group were trained in an auditory fear conditioning paradigm whereby one sound (the conditioned stimulus, or CS^+^) was paired with a foot shock (the unconditioned stimulus, or US), and another sound (the neutral sound, or CS^−^) was not paired with any outcome (Figure 2C; Figure S3A; Methods).

**Figure 2.**
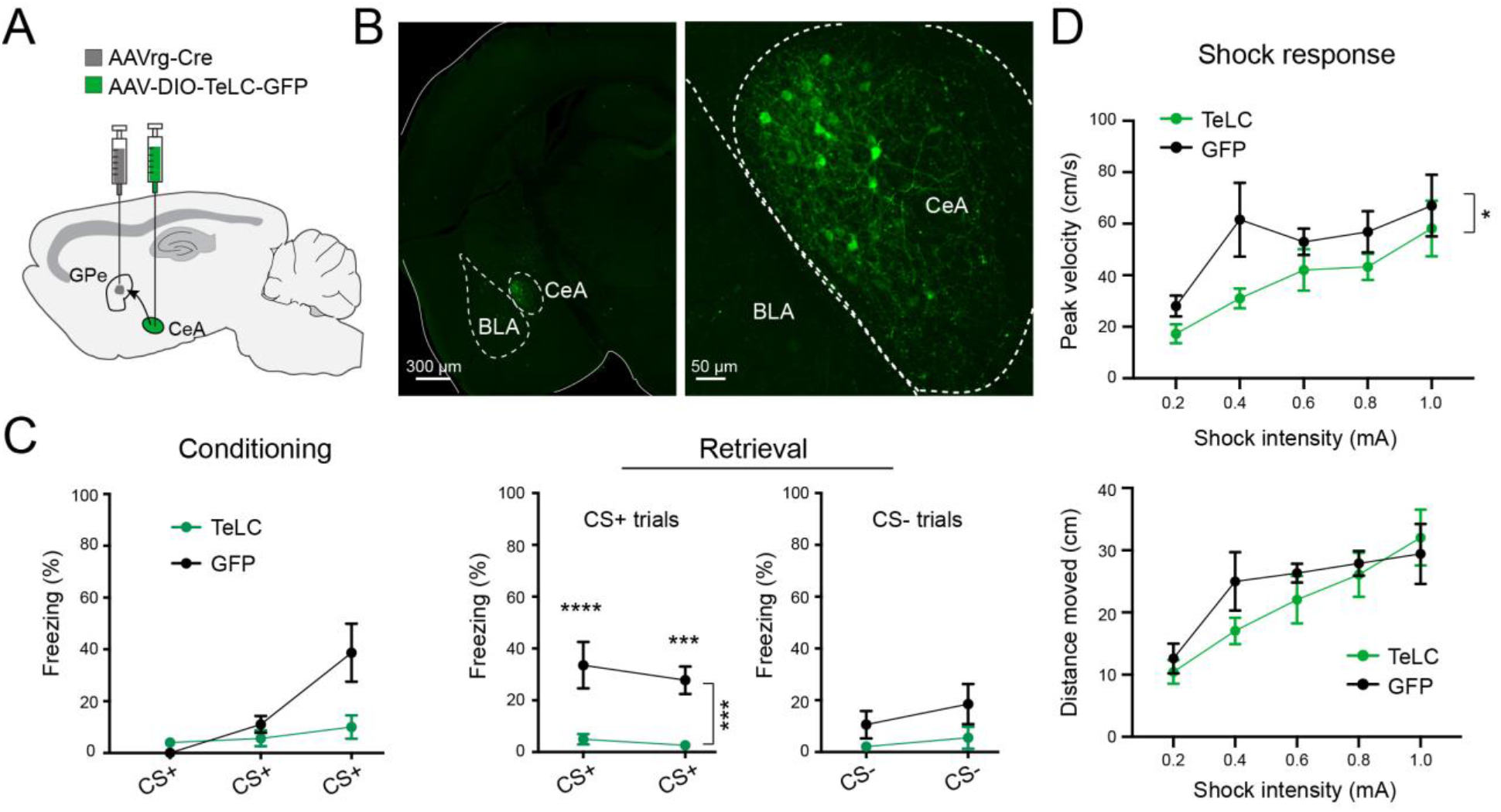
Inhibition of GPe-projecting CeA neurons blocks fear conditioning. (A) A schematic of the approach. (B) Representative confocal images showing the GPe-projecting CeA neurons expressing TeLC. On the right is a higher magnification image of the amygdala area on the left. (C) Freezing behavior in mice in which GPe-projecting CeA neurons expressed TeLC (n = 11) or GFP (n = 6), during Conditioning (left) and Retrieval (right) sessions (conditioning: F(1,15) = 4.47, p = 0.052; retrieval, CS^+^ trials: F(1,15) = 25.21, ***p = 0.0002; ***p < 0.001, ****p < 0.0001; retrieval, CS^−^ trials: F(1,15) = 14.41, p = 0.060; two-way ANOVA with repeated measures, followed by Sidak’s test). (D) Peak velocity (top) and distance moved (bottom) for movements in mice in (C), in response to shocks of varying intensities (peak velocity: F(1,75) = 6.359, *p=0.014; distance moved: F(1,75) = 1.619, p = 0.210; two-way ANOVA). Data in C and D are presented as mean ± s.e.m.

Remarkably, blocking transmitter release from Sst^CeA-GPe^ neurons with TeLC completely abolished the conditioned freezing induced by CS^+^ during a memory retrieval test 24 hours after the conditioning (Figure 2C). Furthermore, this manipulation also reduced the responses of the mice to foot-shocks, as indicated by a reduction in the peak velocity of shock-induced movements (Figure 2D). These results indicate that Sst^CeA-GPe^ neurons are indispensable for fear conditioning, and suggest that these neurons have a role in processing information about the aversive US.

### Sst^CeA-GPe^ neurons represent the unconditioned stimulus during fear conditioning

To further understand the *in vivo* function of Sst^CeA-GPe^ neurons, we recorded the activities of these neurons in behaving mice. For this purpose, we introduced the genetically encoded calcium indicator GCaMP6 (Chen et al., 2013) into these neurons using the above described intersectional viral strategy, in which we injected the AAVrg-Cre unilaterally into the GPe (Figure 1C, D; Figure 2A), and an AAV expressing GCaMP6 in a Cre-dependent manner into the ipsilateral CeA (Figure 3A, B) in wild-type mice. These mice were then implanted with optical fibers above the infected area in the CeA (Figure 3A, B; Figure S4). Four weeks after the surgery, we trained the mice in auditory fear conditioning as described above (Figure 2C), and verified that these mice showed discriminative learning as indicated by higher freezing levels to CS^+^ than to CS^−^ during the memory retrieval test (Figure 3C).

**Figure 3.**
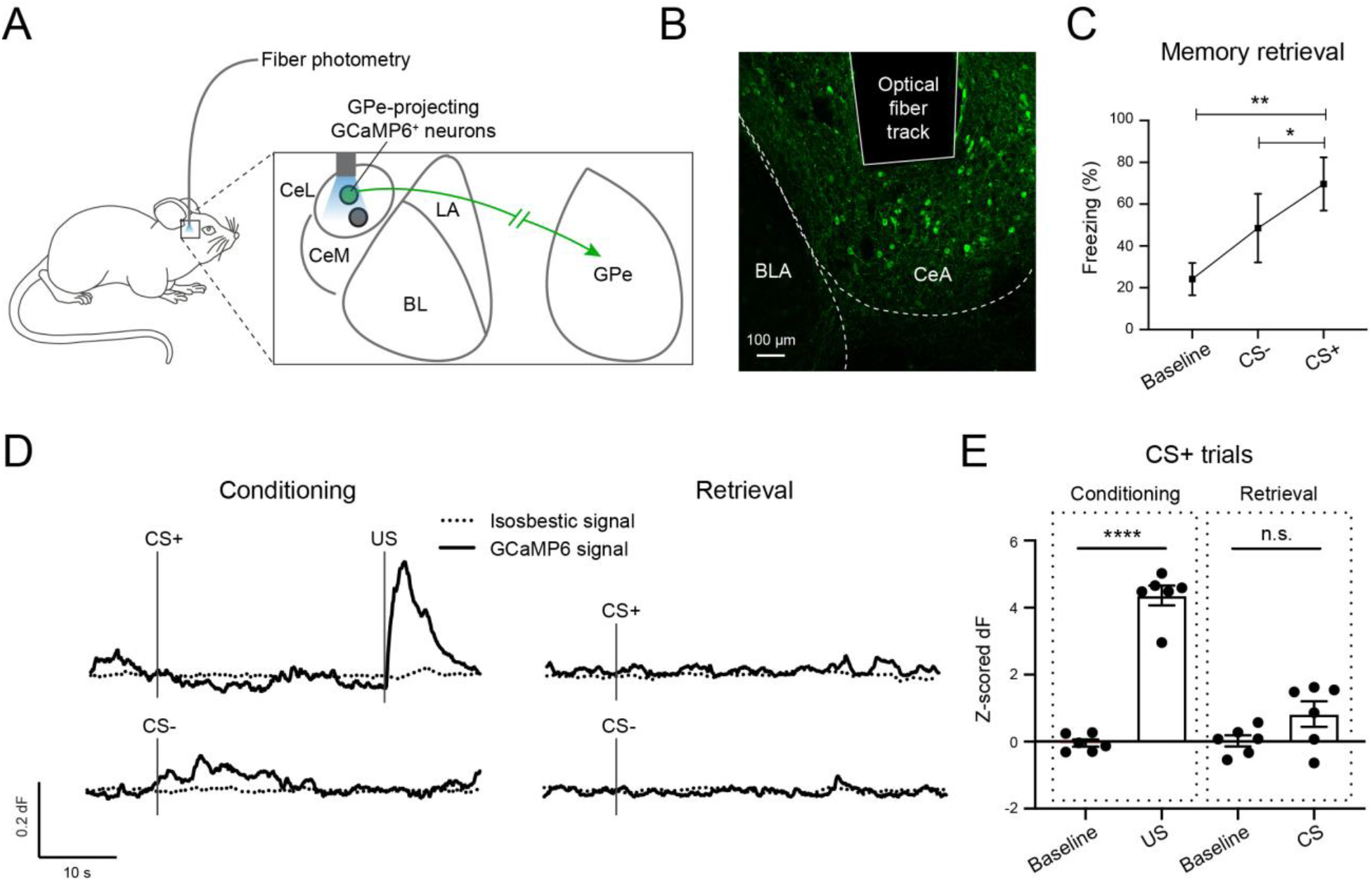
GPe-projecting CeA neurons encode the information about US during fear conditioning. (A) A schematic of the approach. (B) A representative confocal image showing the GPe-projecting CeA neurons expressing GCaMP6f. The track of the implanted optic fiber is also shown. (C) Quantification of freezing behavior during Retrieval (F(1.314, 6.570) = 15.37, p=0.005,*p=0.023, **p=0.005; one-way ANOVA followed by Tukey’s test). (D) Calcium-dependent (solid) and the simultaneously recorded isosbestic (dotted) GCaMP6 fluorescence signals in a representative mouse in CS^+^ and CS^−^ trials for Conditioning (left), and Retrieval (right) sessions. (E) Quantification of the calcium-dependent activities in CS^+^ trials during Conditioning (left) and Retrieval (right) (n = 6 mice; F(3,15) = 80.30, p<0.0001, ****p<0.0001, n.s. (nonsignificant), p>0.05; two-way ANOVA followed by Tukey’s test). Data in C and E are presented as mean ± s.e.m.

We recorded bulk GCaMP6 signals from the infected Sst^CeA-GPe^ neurons in these animals with fiber photometry (Yu et al., 2016) throughout fear conditioning (Figure 3A-D; Figure S4). In this experiment, we simultaneously recorded both the calcium-dependent signals and the isosbestic signals from the GCaMP6 (Figure 3D), with the latter serving to monitor potential motion artifacts (Kim et al., 2016). Notably, we found that Sst^CeA-GPe^ neurons showed potent excitatory response to US (shock) presentations during conditioning, but little response to CS^+^ (or CS^−^) presentations during either conditioning or the memory retrieval test (Figure 3D, E). This result is in sharp contrast with those from Sst^+^ CeA neurons with unknown projection targets, which show robust excitatory responses to CS after fear conditioning as assessed by *in vivo* single unit recording (Fadok et al., 2017) or fiber photometry (Yu et al., 2016). Further examination revealed that the responses of Sst^CeA-GPe^ neurons were significantly higher to stronger shocks than to weaker ones (Figure S5), indicating that the responses represent shock intensity. These results point to the possibility that Sst^CeA-GPe^ neurons play an important role in processing US information thereby instructing learning in fear conditioning.

### Sst^CeA-GPe^ neuron activity during US presentation is required for learning

To determine whether the excitatory response of Sst^CeA-GPe^ neurons evoked by US during fear conditioning is required for learning, we sought to transiently inhibit these neurons only during the presentation of the US. To achieve this goal, we introduced the light sensitive *Guillardia theta* anion-conducting channelrhodopsin 1 (GtACR1) (Govorunova et al., 2015; Mahn et al., 2018) selectively into Sst^CeA-GPe^ neurons using the intersectional viral strategy described above (Figure 1C, D; Figure 2A, B; Figure 3A, B). Specifically, we injected the AAVrg-Cre bilaterally into the GPe and an AAV expressing GtACR1, or GFP, in a Cre-dependent manner bilaterally into the CeA, followed by implanting optical fibers above the infected areas (Figure 4A; Figure S6).

**Figure 4.**
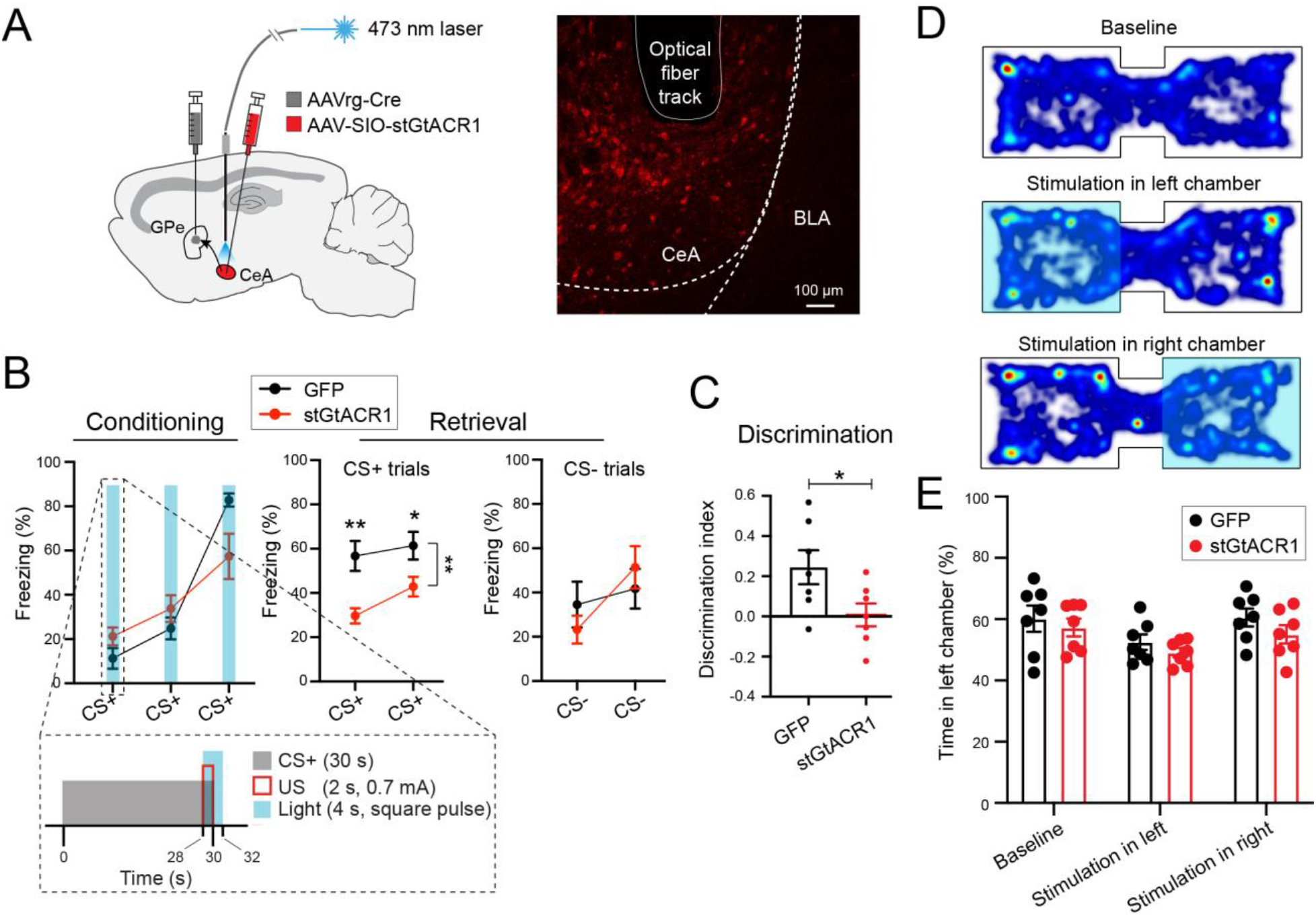
GPe-projecting CeA neuron activity during US presentation is necessary for learning during fear conditioning. (A) Left: a schematic of the approach. Right: a representative confocal image showing the GPe-projecting CeA neurons expressing stGtACR1. The track of the implanted optic fiber is also shown. (B) Freezing behavior in mice in which GPe-projecting CeA neurons expressed stGtACR1 (n = 7) or GFP (n = 7), during Conditioning (left) and Retrieval (right) sessions (conditioning: F(1,12) = 0.117, p > 0.05; retrieval, CS^+^ trials: F(1,12) = 15.65, **p = 0.002; *p < 0.05, **p < 0.010; retrieval, CS^−^ trials: F(1,12) = 0.010, p > 0.05; two-way ANOVA with repeated measures, followed by Sidak’s test). Inset shows the structure and timing of CS^+^, US and light delivery. (C) Discrimination Index calculated as [CS^+^ − CS^−^ / [CS^+^ + CS^−^], where CS^+^ and CS^−^ represent the average freezing during the presentation of CS^+^ and CS^−^, respectively (t(10.51) = 2.329, *p=0.041, Welch’s t-test). (D) Heat-maps for the activity of a representative mouse at baseline (top), or in a situation whereby entering the left (middle) or right (bottom) side of the chamber triggered photo-inactivation of GPe-projecting CeA neurons. (E) Quantification of the mouse activity as shown in (D), for mice in which stGtaCR1 (n = 7) or GFP (n = 7) was introduced into GPe-projecting CeA neurons (F(1, 12) = 2.135, p > 0.05; two-way ANOVA with repeated measures). Data in B, C and E are presented as mean ± s.e.m.

Four weeks following viral injection, both the GtACR1 group and the GFP group (which served as the control) were trained in the auditory fear conditioning paradigm (Figure 4B; Figure S3B). During conditioning, square pulses of blue light, covering the duration of the three US presentations, were delivered to the CeA through the implanted optical fibers (Figure 4B). Notably, we found that this manipulation caused a decrease in CS^+^-induced conditioned freezing behavior in the GtACR1 mice compared with the GFP mice in the retrieval test 24 hours after fear conditioning (Figure 4B). As a result, the ability to discriminate between CS^+^ and CS^−^, quantified as a discrimination index (Methods), was also reduced in the GtACR1 mice (Figure 4C). We next tested these mice in a real-time place preference or aversion (RTPP or RTPA, respectively) task, in which the photo-inhibition was contingent on entering one side of a chamber containing two compartments (Figure 4D). The two groups of animals behaved similarly in this task (Figure 4E), showing no preference or aversion to either side of the chamber. This observation suggests that photo-inhibition of Sst^CeA-GPe^ neurons is not inherently aversive or rewarding. These results indicate that the activities of Sst^CeA-GPe^ neurons during US presentation are required for memory formation in fear conditioning.

### Activation of Sst^CeA-GPe^ neurons during US presentation promotes fear learning

Given that inhibition of Sst^CeA-GPe^ neurons specifically during US presentation impaired learning (Figure 4), it follows that the opposite manipulation, i.e., activation of these neurons specifically during US presentation, might enhance learning in fear conditioning. To test this idea, we introduced ChR2, or GFP, bilaterally into Sst^CeA-GPe^ neurons of wild-type mice using the intersectional viral strategy, followed by optical fiber implantation in the CeA as described above (see Figure 2A, B; Figure 3A, B; Figure 4A; and Figure 5A). We subsequently trained the mice in a mild version of the fear conditioning paradigm (Figure 5B; Figure S3C), in which a weak (0.4 mA) shock was used as the US to avoid the potential ceiling effect a stronger US might have on learning.

**Figure 5.**
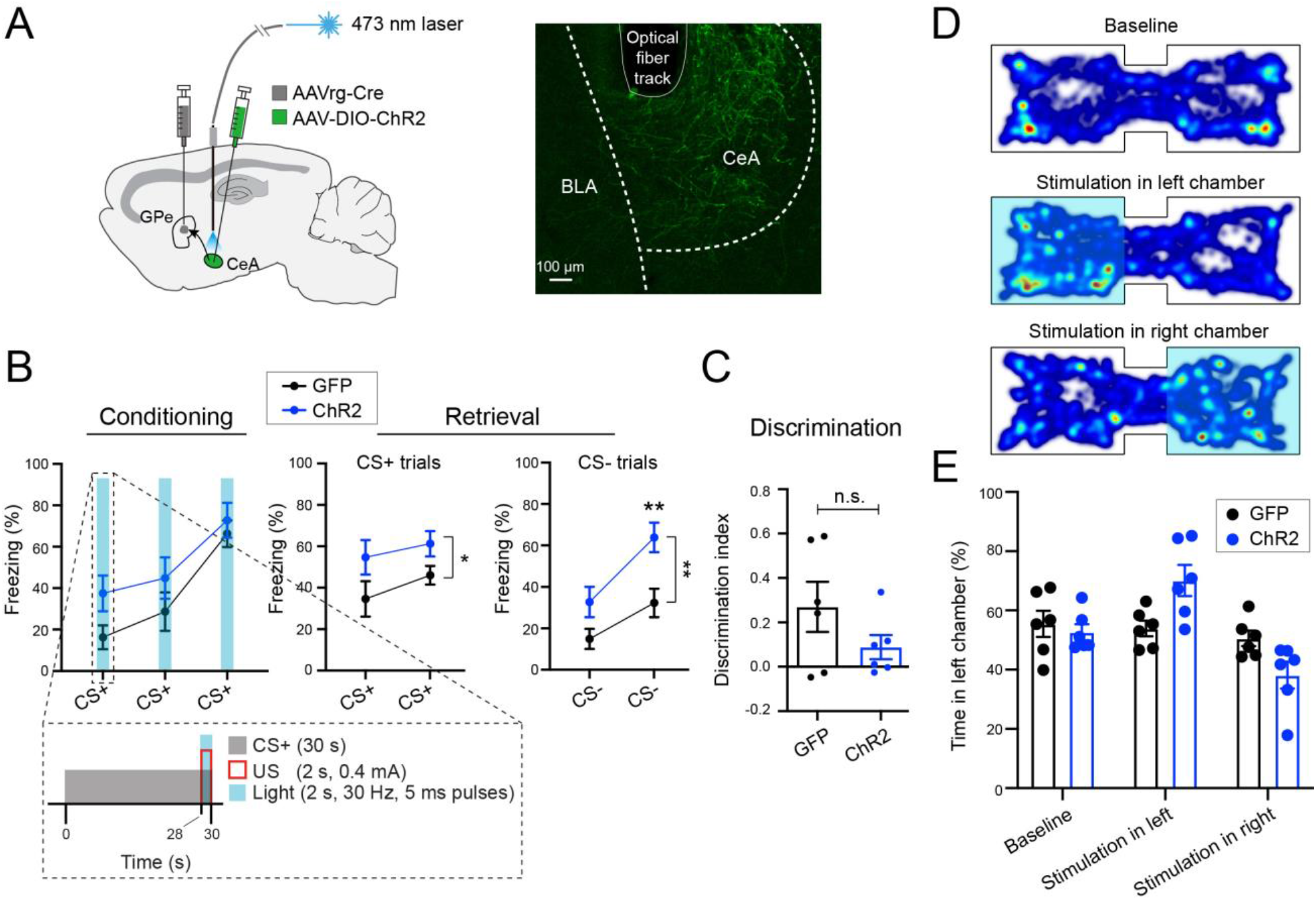
Activation of GPe-projecting CeA neurons during US presentation promotes fear learning. (A) Left: a schematic of the approach. Right: a representative confocal image showing the GPe-projecting CeA neurons expressing ChR2. The track of the implanted optic fiber is also shown. (B) Freezing behavior in mice in which GPe-projecting CeA neurons expressed ChR2 (n = 6) or GFP (n = 6), during Conditioning (left) and Retrieval (right) sessions (conditioning: F(1,10) = 3.682, p=0.084; retrieval, CS^+^ trials: F(1,10) = 5.560, *p = 0.040; retrieval, CS^−^ trials: F(1,10) = 16.34, **p = 0.002; **p < 0.010; two-way ANOVA with repeated measures, followed by Sidak’s test). Inset shows the structure and timing of CS^+^, US and light delivery. (C) Discrimination Index calculated as [CS^+^ − CS^−^ / [CS^+^ + CS^−^], where CS^+^ and CS^−^ represent the average freezing during the presentation of CS^+^ and CS^−^, respectively (t(7.223) = 1.446, p > 0.05, Welch’s t-test). (D) Heat-maps for the activity of a representative mouse at baseline (top), or in a situation whereby entering the left (middle) or right (bottom) side of the chamber triggered photo-activation of GPe-projecting CeA neurons. (E) Quantification of the mouse activity as shown in (D), for mice in which ChR2 (n = 6) or GFP (n = 6) was introduced into GPe-projecting CeA neurons (F(1,10) = 0.019, p > 0.05; two-way ANOVA with repeated measures). Data in B, C and E are presented as mean ± s.e.m.

During conditioning, three brief trains of photo-stimulation, each coinciding with a US presentation, were delivered to the CeA (Figure 5B). This manipulation increased CS^+^-induced conditioned freezing behavior in the ChR2 mice compared with the GFP mice in a retrieval test 24 hours after the conditioning (Figure 5B). Interestingly, the ChR2 mice also showed an increase in freezing response to CS^−^ during the retrieval test (Figure 5B), albeit their discrimination index did not significantly differ from that of the GFP mice (*P* = 0.19, Welch’s t-test; Figure 5C). To check if the facilitating effect on learning is because activating Sst^CeA-GPe^ neurons influences valence processing, we tested these mice again in the RTPP or RTPA task for photo-stimulating Sst^CeA-GPe^ neurons using the same parameters as those used in fear conditioning. Notably, the two groups of animals behaved similarly in the test (Figure 5D, E), indicating that photo-activation of Sst^CeA-GPe^ neurons is not inherently aversive or rewarding. These results together suggest that activating Sst^CeA-GPe^ neurons during US presentation promotes the formation of fear memories, although the activation may not by itself produce aversive valence.

## Discussion

Animals have the ability to use an environmental cue (i.e., CS) to predict the occurrence of an aversive or harmful consequence (i.e., US) – on condition that the former is frequently associated with the latter – and to show appropriate behavioral reactions based on the prediction (Lang and Davis, 2006; LeDoux, 2000; Pavlov, 1927; Schultz, 2006). Such ability is fundamental for survival and adaptation to the environment. Extensive studies, exemplified by those focusing on Pavlovian fear conditioning, have shown that the CeA plays important roles in the establishment of adaptive defensive behaviors (Duvarci and Pare, 2014; Fadok et al., 2018; Herry and Johansen, 2014; Janak and Tye, 2015; Li, 2019). However, despite the intensive study, how the CeA processes and represents the aversive US during fear conditioning, and how it contributes to the formation of aversive memories remain to be fully understood. Here, we identified a previously unknown circuit, the Sst^CeA-GPe^ circuit, that is essential for fear conditioning. Specifically, we showed that Sst^+^ CeA neurons send a major projection to innervate GPe neurons, and permanent inhibition of Sst^CeA-GPe^ neurons prevented fear conditioning. Moreover, Sst^CeA-GPe^ neurons were excited by US but not CS during fear conditioning, and transient inactivation or activation of these neurons specifically during US presentation impaired or promoted, respectively, fear learning. One the basis of these results, we propose that the major function of Sst^CeA-GPe^ neurons in fear conditioning is to represent and process the US information, and convey this information to downstream GPe neurons, thereby instructing learning.

The GPe is a major basal ganglia structure whose roles in motor control have been the focus of investigation (Kita, 2007; Wallace et al., 2017), but whose other functions have been understudied. Nevertheless, the GPe has been implicated in regulating emotions or affects, including fear or threat. For example, human imaging studies indicate that GPe activation is associated with negative emotions, such as fear, disgust, depression and anxiety (Binelli et al., 2014; Hattingh et al., 2012; Ipser et al., 2013; Murphy et al., 2003). In addition, animal studies have shown that lesions and pharmacological or molecular manipulations in the GPe potently alter fear- or anxiety-like behaviors (Blanchard et al., 1981; Hernadi et al., 1997; Kertes et al., 2009; Sztainberg et al., 2011; Talalaenko et al., 2006). These findings thus ascribe a function of fear or threat regulation to the GPe. An obvious question is how this GPe function is related to that of the known “fear circuit”, including the amygdala. A potential anatomical link between the GPe and the fear circuit is suggested by previous studies, which demonstrate the existence of the CeA to GPe projections (Hunt et al., 2018; Shinonaga et al., 1992). Nevertheless, the roles of these projections in fear regulation, and in behavior in general, have remained unknown.

Our study uncovers that these projections originate mainly from Sst^+^ CeA neurons and shows that the Sst^CeA-GPe^ circuit indeed constitutes a neural substrate for fear learning. The activities of Sst^CeA-GPe^ neurons may not be sufficient to cause aversive responses, as suggested by the observation that activating these neurons produced no effect in the RTPP/RTPA test. However, the information carried by these neurons could be important for valence processing in the GPe. Future studies need to elucidate how GPe neurons interact with the upstream Sst^CeA-GPe^ neurons and neurons in downstream structures to participate in fear processing and learning.

Sst^+^ CeA neurons send long-range projections to a number of target areas (Ahrens et al., 2018; Fadok et al., 2018; Li, 2019; Penzo et al., 2014; Steinberg et al., 2020; Ye and Veinante, 2019; Yu et al., 2017; Zhou et al., 2018). Some of these projections have been studied in the context of fear conditioning or anxiety-related behaviors (Ahrens et al., 2018; Penzo et al., 2014; Steinberg et al., 2020; Zhou et al., 2018). However, the encoding properties of these projections and how they contribute to specific aspects of learning or executing defensive behaviors have not been characterized. Our study pinpoints the main function of Sst^CeA-GPe^ neurons being representation and processing of US information during fear conditioning. Future studies need to delineate whether and how different CeA projection pathways differentially but coordinately contribute to the establishment of defensive behaviors.

## Materials and Methods

### Animals

Male and female mice of 3-6 months old were used in the behavioral experiments; those of 6-10 weeks old were used in the *in vitro* electrophysiology experiments. Mice were housed under a 12-h light/dark cycle (7 a.m. to 7 p.m. light) in groups of 2-5 animals, with food and water available *ad libitum*. All behavioral experiments were performed during the light cycle.

Littermates were randomly assigned to different groups prior to experiments. All mice were bred onto a C57BL/6J background. All experimental procedures were approved by the Institutional Animal Care and Use Committee of Cold Spring Harbor Laboratory (CSHL) and performed in accordance to the US National Institutes of Health guidelines.

The C57/B6 wild-type mice were purchased from the Jackson Laboratory. The *H2B-GFP* (*Rosa26-stop*^*flox*^*-H2B-GFP*) reporter mouse line (He et al., 2012) was generated by Z. Josh Huang’s lab at CSHL. The *Sst-IRES-Cre* mice (Taniguchi et al., 2011) were purchased from the Jackson Laboratory (Stock No: 013044). The *Ai14* reporter mice (Madisen et al., 2010) were purchased from the Jackson Laboratory (Stock No: 007908).

### Viral vectors and reagents

The retrograde AAV expressing Cre (AAVrg-Cre), which is suitable for retrogradely labeling CeA neurons, was newly developed and packed in Xiaoke Chen’s lab at Stanford University. The AAV2/9-CAG-DIO-TeLC-eGFP was previously described (Murray et al., 2011) and custom-packed at Penn Vector Core (Philadelphia, PA, USA). The AAV9-EF1a-DIO-hChR2(H134R)-eYFP-WPRE-hGH were made by Penn Vector Core. The AAV9-CAG-Flex-GFP was produced by the University of North Carolina vector core facility (Chapel Hill, North Carolina, USA). The AAV1.Syn.Flex.GCaMP6f.WPRE.SV40, AAV1-hSyn1-SIO-stGtACR1-FusionRed and AAV2-hSyn-DIO-mCherry were produced by Addgene (Watertown, MA, USA). All viral vectors were stored in aliquots at −80°C until use.

The retrograde tracer cholera toxin subunit B (CTB) conjugated with either Alexa Fluor™ 647 or 555 (CTB-647 or CTB-555, respectively) was purchased from Invitrogen, Thermo Fisher Scientific (Waltham, Massachusetts, USA). CTB was used at a concentration of 1mg/ml in phosphate-buffered saline.

### Stereotaxic Surgery

Standard surgical procedures were followed for stereotaxic injection (Li et al., 2013; Penzo et al., 2015; Yu et al., 2017; Yu et al., 2016). Briefly, mice were anesthetized with isoflurane (3% at the beginning and 1% for the rest of the surgical procedure), and positioned in a stereotaxic injection frame (myNeuroLab.com). A digital mouse brain atlas was linked to the injection frame to guide the identification and targeting (Angle Two Stereotaxic System, myNeuroLab.com). The injection was performed at the following stereotaxic coordinates for CeL: −1.22 mm from Bregma, 2.9 mm lateral from the midline, and 4.6 mm vertical from skull surface; for GPe: −0.46 mm from Bregma, 1.85 mm lateral from the midline, and 3.79 mm vertical from skull surface; and for BNST: 0.20 mm from bregma, 0.85 mm lateral from the midline, and 4.15 mm vertical from skull surface.

For virus or tracer injection, we made a small cranial window (1–2 mm^2^), through which virus or fluorescent tracers (~0.3 μl) were delivered via a glass micropipette (tip diameter, ~5 μm) by pressure application (5–20 psi, 5–20 ms at 0.5 Hz) controlled by a Picrospritzer III (General Valve) and a pulse generator (Agilent). During the surgical procedure, mice were kept on a heating pad maintained at 35°C and were brought back to their home-cage for post-surgery recovery and monitoring. Subcutaneous Metacam (1-2 mg kg–1 meloxicam; Boehringer Ingelheim Vetmedica, Inc.) was given post-operatively for analgesia and anti-inflammatory purposes. For optogenetic experiments, optical fibers (200 μm diameter, 0.22 NA, 5 mm length) were implanted bilaterally 0.3 mm over the CeA. A small metal bar, which was used to hold the mouse in the head fixation frame to connect optical fibers during training, was mounted on the skull with C&B Metabond quick adhesive cement (Parkell Inc.), followed by dental cement (Lang Dental Manufacturing Co., Inc.).

### *In vitro* electrophysiology

For the *in vitro* electrophysiology experiments, mice were anaesthetized with isoflurane and perfused intracardially with 20 mL ice-cold artificial cerebrospinal fluid (ACSF) (118 mM NaCl, 2.5 mM KCl, 26.2 mM NaHCO_3_, 1 mM NaH_2_PO_4_, 20 mM glucose, 2 mM MgCl_2_ and 2 mM CaCl_2_, pH 7.4, gassed with 95% O_2_ and 5% CO_2_). Mice were then decapitated and their brains quickly removed and submerged in ice-cold dissection buffer (110.0 mM choline chloride, 25.0 mM NaHCO_3_, 1.25 mM NaH_2_PO_4_, 2.5 mM KCl, 0.5 mM CaCl_2_, 7.0 mM MgCl_2_, 25.0 mM glucose, 11.6 mM ascorbic acid and 3.1mM pyruvic acid, gassed with 95% O_2_ and 5% CO_2_). 300 μm coronal slices containing the globus pallidus externa (GPe) were cut in dissection buffer using a HM650 Vibrating-blade Microtome (Thermo Fisher Scientific). Slices were immediately transferred to a storage chamber containing ACSF at 34 °C. After 40 min recovery time, slices were transferred to room temperature (20–24°C) and perfused with gassed ACSF constantly throughout recording.

Whole-cell patch clamp recording was performed as previously described (Li et al., 2013). Briefly, recording from GPe neurons was obtained with Multiclamp 700B amplifiers and pCLAMP 10 software (Molecular Devices, Sunnyvale, California, USA), and was visually guided using an Olympus BX51 microscope equipped with both transmitted and epifluorescence light sources (Olympus Corporation, Shinjuku, Tokyo, Japan). The external solution was ACSF. The internal solution contained 115 mM cesium methanesulfonate, 20 mM CsCl, 10 mM HEPES, 2.5 mM MgCl_2_, 4 mM Na_2_ATP, 0.4 mM Na_3_GTP, 10 mM sodium phosphocreatine and 0.6 mM EGTA (pH 7.2).

As the acute slices were prepared from *Sst-IRES-Cre* mice in which Sst^+^ CeA neurons were infected with AAV expressing ChR2-YFP, to evoke synaptic transmission onto GPe neurons driven by Sst^CeA-GPe^ neurons, a blue light was used to stimulate ChR2-expressing axons originating from Sst^CeA-GPe^ neurons. The light source was a single-wavelength LED system (λ = 470 nm; http://www.coolled.com/) connected to the epifluorescence port of the Olympus BX51 microscope. A light pulse of 1 ms, triggered by a TTL signal from the Clampex software, was delivered every 10 seconds to evoke synaptic responses. Evoked inhibitory post-synaptic currents (IPSCs) were recorded at a holding potential of 0 mV and in ACSF with 100 μM AP5 and 10 μM CNQX added to block excitatory synaptic transmission. Synaptic responses were low-pass filtered at 1 kHz and were analyzed using pCLAMP 10 software. Evoked IPSCs were quantified as the mean current amplitude from 50-60 ms after stimulation.

### Immunohistochemistry

For histology analysis, mice were anesthetized with Euthasol (0.2 mL; Virbac, Fort Worth, Texas, USA) and perfused transcardially with 30 mL cold phosphate buffered saline (PBS) followed by 30 mL 4% paraformaldehyde (PFA) in PBS. Brains were removed immediately from the skull and placed in PFA for at least 24 hours and then in 30% sucrose in PBS solution for 24 hours for cryoprotection. Coronal sections (50 μm) were cut using a freezing microtome (Leica SM 2010R, Leica) and placed in PBS in 12-well plates. Brain sections were first washed in PBS (3 × 5 min), incubated in PBST (0.3% Triton X-100 in PBS) for 30 min at room temperature (RT) and then washed with PBS (3 × 5 min). Next, sections were blocked in 5% normal goat serum in PBST for 30 min at RT and then incubated with the primary antibody for 12 h at 4 °C. Sections were washed with PBS (5 × 15 min) and incubated with the fluorescent secondary antibody at RT for 2 h. After washing with PBS (5 × 15 min), sections were mounted onto slides with Fluoromount-G (eBioscience, San Diego, California, USA). Images were taken using an LSM 710 laser-scanning confocal microscope (Carl Zeiss, Oberkochen, Germany).

The primary antibodies used in this study were: chicken anti-GFP (Aves Labs, catalogue number GFP1020, lot number GFP697986), rabbit anti-RFP (Rockland, catalogue number 600-401-379, lot number 34135). The fluorophore-conjugated secondary antibodies used were Alexa Fluor® 488 donkey anti-chicken IgG (H+L), Alexa Fluor® 488 goat anti-rabbit IgG (H+L) and Alexa Fluor® 555 goat anti-rabbit IgG (H+L) (Life Technologies, Carlsbad, California, USA).

### Fluorescent *in situ* hybridization

Single molecule fluorescent *in situ* hybridization (smFISH) (ACDBio, RNAscope) was used to detect the expression of *Sst* and *Prkcd* mRNAs in the central amygdala (CeA) of adult mice, which were injected in the GPe with CTB-555. 5 days after CTB injection, mice were first anesthetized under isoflurane and then decapitated. Their brain tissue was first embedded in cryomolds (Sakura Finetek, Ref 4566) filled with M-1 Embedding Matrix (Thermo Scientific, Cat. No. 1310) then quickly fresh-frozen on dry ice. The tissue was stored at −80 °C until it was sectioned with a cryostat. Cryostat-cut sections (16-μm) containing the CeA were collected and quickly stored at −80 °C until processed. Hybridization was carried out using the RNAscope kit (ACDBio).

The day of the experiment, frozen sections were post-fixed in 4% PFA in RNA-free PBS (hereafter referred to as PBS) at RT for 15 min, then washed in PBS, dehydrated using increasing concentrations of ethanol in water (50%, once; 70%, once; 100%, twice; 5 min each). Sections were then dried at RT and incubated with Protease IV for 30 min at RT. Sections were washed in PBS three times (5 min each) at RT, then hybridized. Probes against *Sst* (Cat. No. # 404631, dilution 1:50) and *Prkcd* (Cat. No. # 441791, dilution 1:50) were applied to CeA sections. Hybridization was carried out for 2 h at 40°C. After that, sections were washed twice in PBS (2 min each) at RT, then incubated with three consecutive rounds of amplification reagents (30 min, 15 min and 30 min, at 40°C). After each amplification step, sections were washed twice in PBS (2 min each) at RT. Finally, fluorescence detection was carried out for 15 min at 40°C. The red channel was left free for detection of CTB-555 fluorescence. Sections were then washed twice in PBS, incubated with DAPI for 2 min, washed twice in PBS (2 min each), then mounted with coverslip using mounting medium. Images were acquired using an LSM780 confocal microscope equipped with 20x, 40x or 63x lenses, and visualized and processed using ImageJ and Adobe Illustrator.

### Behavioral tasks

#### Auditory fear conditioning

We followed standard procedures for conventional auditory fear conditioning (Li et al., 2013; Penzo et al., 2014; Penzo et al., 2015; Yu et al., 2017). Briefly, mice were initially handled and habituated to a conditioning cage, which was a Mouse Test Cage (18 cm × 18 cm × 30 cm) with an electrifiable floor connected to a H13-15 shock generator (Coulbourn Instruments, Whitehall, PA). The Test Cage was placed inside a sound attenuated cabinet (H10-24A; Coulbourn Instruments). Before each habituation and conditioning session, the Test Cage was wiped with 70% ethanol. The cabinet was illuminated with white light during habituation and conditioning sessions.

During habituation, two 4-kHz 60-dB tones and two 12-kHz 60-dB tones, each of which was 30 s in duration, were delivered at variable intervals within an 8-minute session. During conditioning, mice received three presentations of the 4-kHz tone (conditioned stimulus; CS^+^), each of which co-terminated with a 2-s 0.7-mA foot shock (unless otherwise stated), and three presentations of the 12-kHz tone, which were not paired with foot shocks (CS^−^). The CS^+^ and CS^−^ were interleaved pseudo-randomly, with variable intervals between 30 and 90 s within a 10-minute session. The test for fear memory (retrieval) was performed 24 h following conditioning in a novel context, where mice were exposed to two presentations of CS^+^ and CS^−^ (>120 s inter-CS interval). The novel context was a cage with a different shape (22 cm × 22 cm × 21 cm) and floor texture compared with the conditioning cage, and was illuminated with infrared light. Prior to each use the floor and walls of the cage were wiped clean with 0.5% acetic acid to make the scent distinct from that of the conditioning cage.

For optogenetic manipulation with stGtACR1 during fear conditioning, blue light (473 nm, 5 mW; 4-s square pulse) was delivered via tethered patchcord to the implanted optical fibers. The onset of the light coincided with the onset of US (2-s 0.7 mA foot shock) presentation. For optogenetic manipulation with ChR2 during fear conditioning, blue light (473 nm, 5 mW; 30-Hz, 5-ms pulses for 2 s) was delivered via tethered patchcord to the implanted optical fibers, coinciding with the presentation of US (2-s 0.4 mA foot shock).

Animal behavior was videotaped with a monochrome CCD-camera (Panasonic WV-BP334) at 3.7 Hz and stored on a personal computer. The FreezeFrame software (Coulbourn Instruments) was used to control the delivery of both tones and foot shocks. Freezing behavior was analyzed with FreezeFrame software (Coulbourn Instruments) for the TeLC experiment. For subsequent fiber photometry and optogenetic experiments, Ethovision XT 5.1 (Noldus Information Technologies) was used to track the animal, and freezing was calculated using a custom Matlab script for improved tracking while avoiding the influence by patchcords and optic fibers attached to animal’s head. Baseline freezing levels were calculated as the average freezing during the first 100 s of the session before any stimuli were presented, and freezing to the auditory stimuli was calculated as the average freezing during the tone presentation. The average of the freezing responses to two CS^+^ or CS^−^ presentations during recall was used as an index of fear. Discrimination Index was calculated as the difference between freezing to the CS^+^ and CS^−^, normalized by the sum of freezing to both tones.

#### Real-time place preference or aversion test

Freely moving mice were habituated to a two-sided chamber (made from Plexiglas; 23 × 33 × 25 cm for each side) for 10 min, during which baseline preference to each side was assessed. During the first test session (10 min), one side of the chamber was designated the photo-stimulation side, and mice were placed in the middle to start the experiment. Once the mouse entered the stimulation side, photo-stimulation (5-ms pulses, 30 Hz, 10 mW (measured at the tip of optic fibers)) with a 473-nm laser (OEM Laser Systems Inc., Bluffdale, Utah, USA) was turned on, and was turned off upon the mouse exiting the stimulation side. In the second test session (10 min) this procedure was repeated, with the opposite side being the stimulation side. Animal behavior was videotaped with a CCD camera (C930, Logitech) and tracked with Ethovision, which was also used to control the laser stimulation and extract behavioral parameters (position, time, distance and velocity).

### *In vivo* fiber photometry and data analysis

A commercial fiber photometry system (Neurophotometrics Ltd., San Diego, CA, USA) was used to record GCaMP6f signals in Sst^CeA-GPe^ neurons *in vivo* in behaving animals through an optical fiber (200 μm fiber core diameter, 5.0 mm length, 0.37 NA; Inper, Hangzhou, China) implanted in the CeA. A patch cord (fiber core diameter, 200 μm; Doric Lenses) was used to connect the photometry system with the implanted optical fiber. The intensity of the blue light (λ = 470 nm) for excitation was adjusted to ~20 μW at the tip of the patch cord. Emitted GCaMP6f fluorescence was bandpass filtered and focused on the sensor of a CCD camera. Photometry signals and behavioral events were aligned based on an analogue TTL signal generated by a Bpod. Mean values of signals from a region of interest were calculated and saved using Bonsai software (Bonsai), and exported to MATLAB for further analysis.

To correct for slow baseline drifting caused by photobleaching, a time-dependent baseline F0(t) was computed as described previously (Jia et al., 2011). The percentage ΔF/F was calculated as 100 × (F(t) − F_0_(t))/F_0_(t), where F(t) is the raw fluorescence signal at time t. After baseline drift correction, the fluorescence signals were z-scored relative to the mean and standard deviation of the signals in a 2 s time window immediately prior to CS onset. In this experiment, we simultaneously recorded both the calcium-dependent signals and the isosbestic signals from the GCaMP6, with the latter serving to monitor potential motion artifacts as previously described (Kim et al., 2016).

### Data Analysis and Statistics

All statistics are indicated where used. Statistical analyses were performed with GraphPad Prism Software (GraphPad Software, Inc., La Jolla, CA). Normality was tested by D’Agostino-Pearson or Shapiro-Wilk normality tests. All behavioral experiments were controlled by computer systems, and data were collected and analyzed in an automated and unbiased way. Virus-injected animals in which the injection site was incorrect were excluded. No other mice or data points were excluded.

## Acknowledgements

We thank members of the Li laboratory for helpful discussions, and Z. Josh Huang for providing the *H2B-GFP* (*Rosa26-stop*^*flox*^*-H2B-GFP*) reporter mice. This work was supported by grants from EMBO (ALTF 458-2017, A.F.), Swedish Research Council (2017-00333, A.F.), Charles H. Revson Senior Fellowships in Biomedical Science (A.F.), the National Institutes of Health (NIH) (R01MH101214, R01MH108924, R01NS104944, B.L.), Human Frontier Science Program (RGP0015/2016, B.L.), the Stanley Family Foundation (B.L.), Simons Foundation (344904, B.L.), Wodecroft Foundation (B.L.), the Cold Spring Harbor Laboratory and Northwell Health Affiliation (B.L.) and Feil Family Neuroscience Endowment (B.L.).

## Author contributions

J.G. and B.L. conceived and designed the study. J.G. conducted the experiments and analyzed data. K.Y. identified the Sst^CeA-GPe^ projections and assisted with experiments. A.F. designed and performed the retrograde tracing combined with smFISH experiments and analyzed data. G.T.N. and X.C. developed the new AAVrg-Cre virus. R.S. assisted with the smFISH experiments. J.G. and B.L. wrote the paper with inputs from all authors.

