## Supplemental Material for "A central amygdala-globus pallidus circuit conveys unconditioned stimulus information and controls fear learning"

### FIGURES AND SUPPLEMENTARY FIGURES

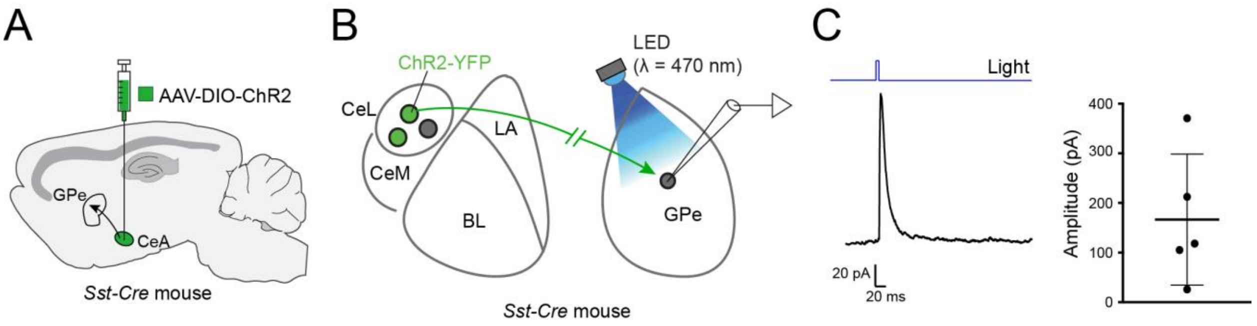


**Figure S1. Functional connectivity between $Sst^{CeA\text{-}GPe}$ neurons and GPe neurons**

(A) A schematic of the approach.

(B) A schematic of the *in vitro* recording configuration in acute slices.

(C) Left: an average trace of synaptic currents recorded from a representative neuron in the GPe. Synaptic currents were evoked by optogenetic activation of axon fibers originating from $Sst^{CeA\text{-}GPe}$ neurons, and were recorded at a holding potential of 0 mV in the presence of 100 μM AP5 and 10 μM CNQX to isolate inhibitory postsynaptic currents (IPSCs; see Methods). The upward square pulse in the blue trace on top of the synaptic current indicates the timing of photo-stimulation. Right: quantification of the amplitude of the evoked IPSCs (5 out of 12 cells recorded from 5 mice had evoked IPSCs).

Data in C is presented as mean ± s.e.m.

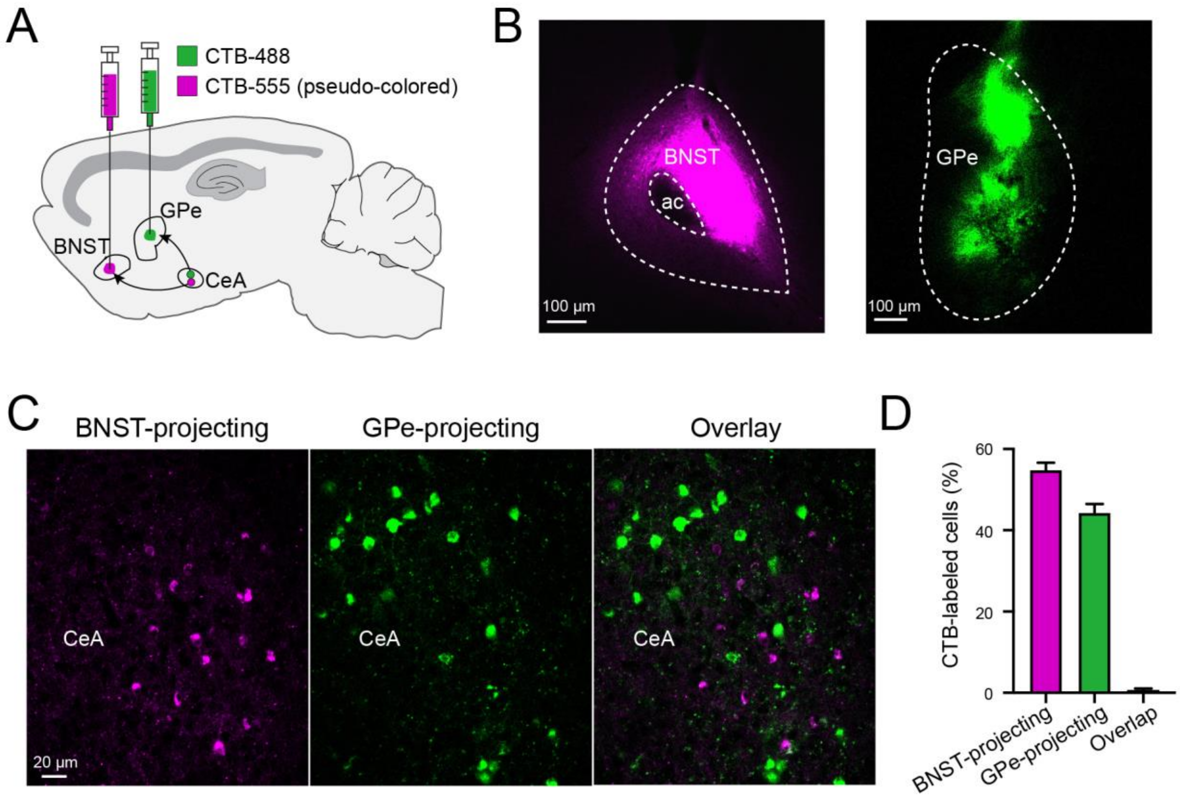


**Figure S2. GPe-projecting CeA neurons do not send collateral projections to the BNST**

(A) A schematic of the approach.

(B) Histology images showing CTB-555 (pseudo-colored) and CTB-488 injection locations in the BNST and GPe, respectively, of a representative mouse.

(C) Representative confocal images of the CeA in the same mouse as that in (B), showing CeA neurons labelled by CTB-555 and CTB-488.

(D) Quantification of the CeA neurons projecting to the BNST, GPe or both structures (n = 2 mice).

Data in D is presented as mean ± s.e.m.

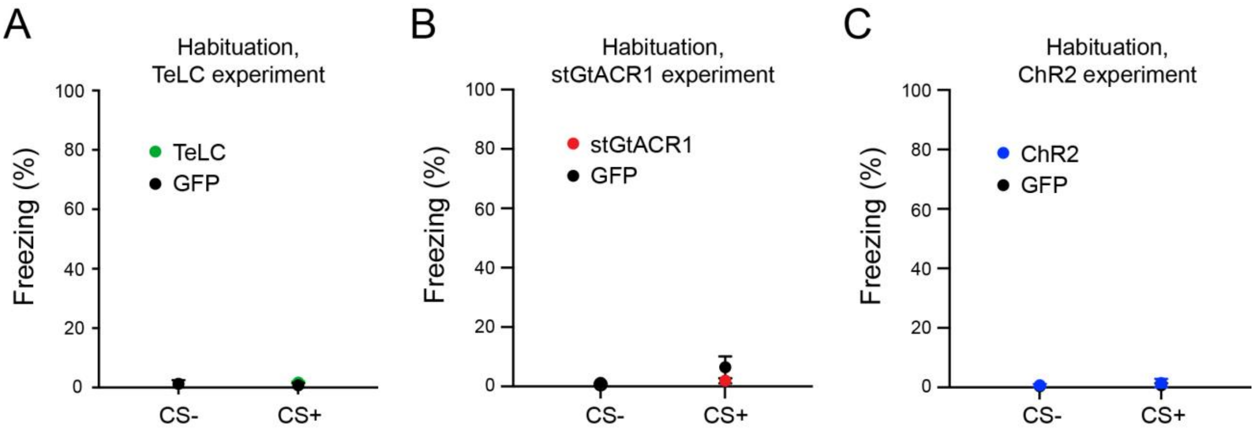


**Figure S3. Freezing behavior during the habituation session in different experiments**
Average freezing to $CS^-$ and $CS^+$ presentations during the habituation session for the TeLC experiment in Figure 2 (A), the stGtACR1 experiment in Figure 4 (B) and ChR2 experiment in Figure 5 (C).

Data are presented as mean ± s.e.m.

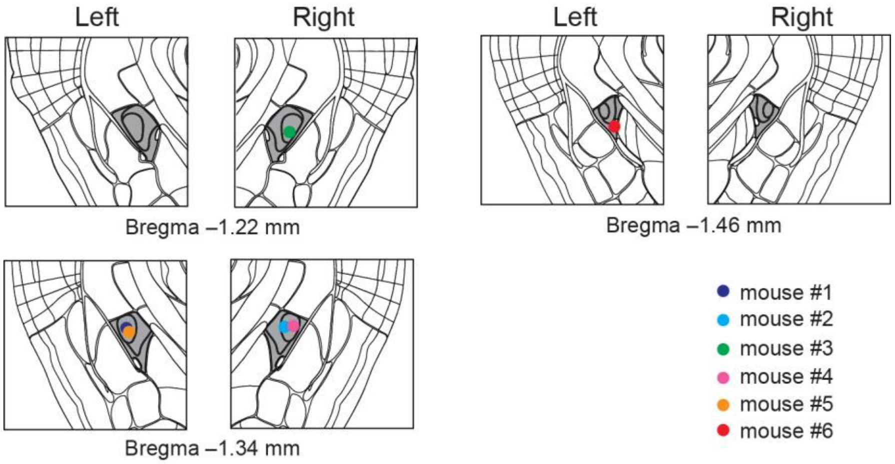


**Figure S4. Optic fiber implant locations for recording Sst$^{CeA\text{-}GPe}$ neuron activities with fiber photometry (related to Figure 3)**

Schematics showing the placement of optic fibers in all the mice used for recording the activities in Sst$^{CeA\text{-}GPe}$ neurons with fiber photometry (n = 6 mice). The CeA is colored in dark gray.

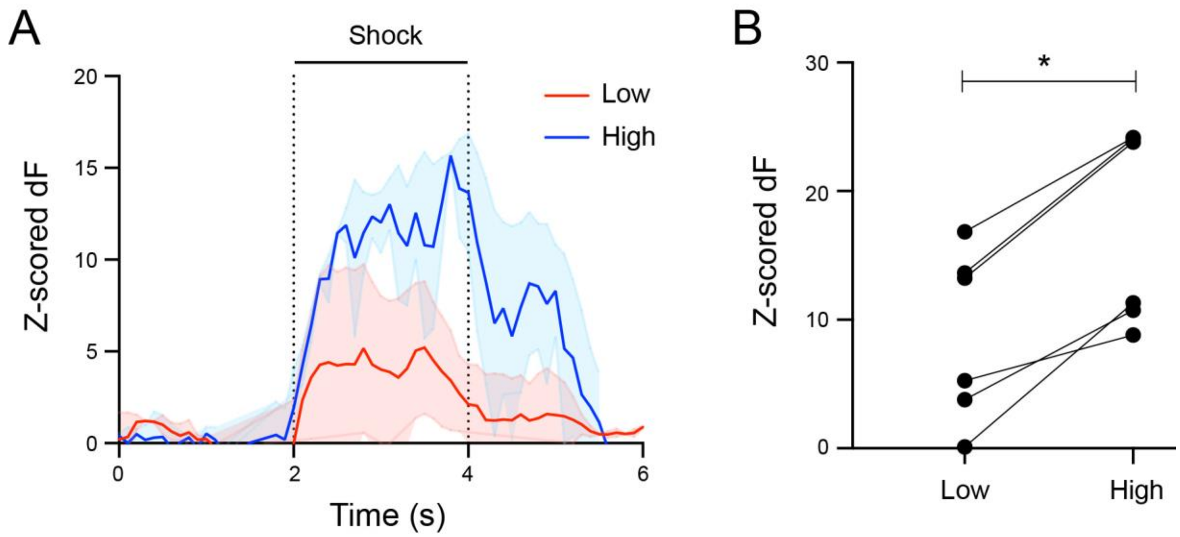


**Figure S5. Sst$^{CeA\text{-}GPe}$ neuron responses scale with shock intensities**

(A) Average responses of Sst$^{CeA\text{-}GPe}$ neurons in an example mouse to shocks of low (0.2 or 0.4 mA) and high (0.8 or 1.0 mA) intensities.

(B) Quantification of responses of all mice (n = 6) to shocks of low and high intensities (*p=0.031, Wilcoxon paired t-test).

Data are presented as mean ± s.e.m.

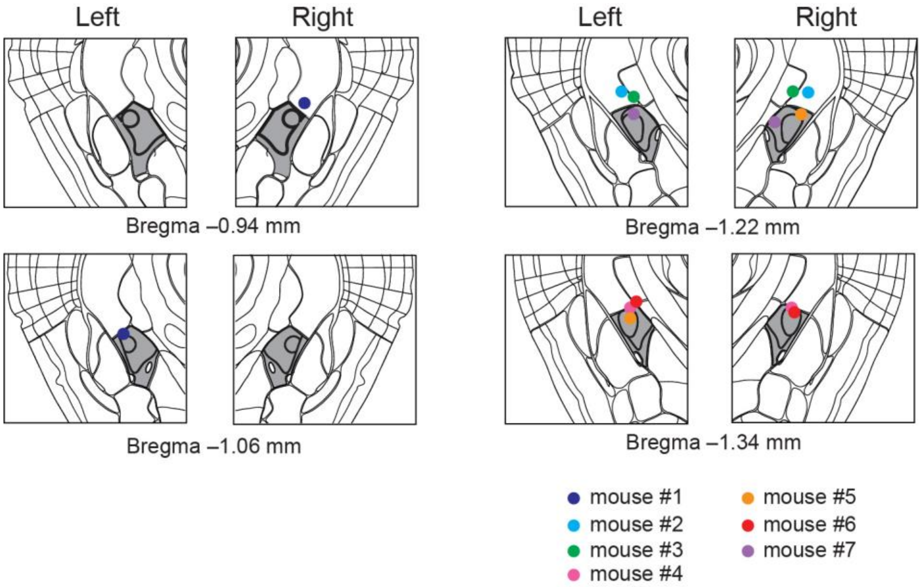


**Figure S6. Optic fiber implant locations for optogenetic inhibition of $Sst^{CeA\text{-}GPe}$ neurons (related to Figure 4)**
Schematics showing the placement of optic fibers in all the mice used for inhibiting $Sst^{CeA\text{-}GPe}$ neurons with optogenetics (n = 7 mice). The CeA is colored in dark gray.

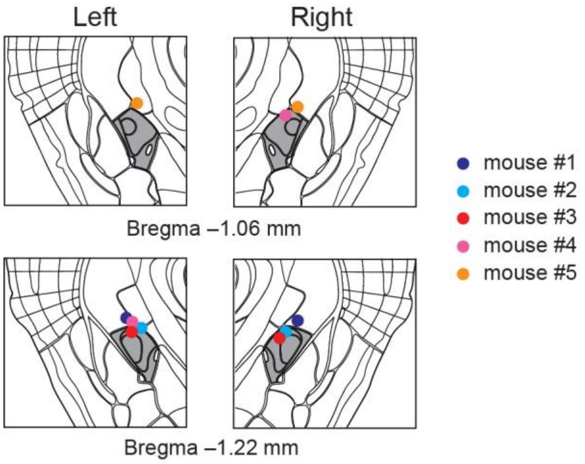


**Figure S7. Optic fiber implant locations for optogenetic activation of Sst$^{CeA\text{-}GPe}$ neurons (related to Figure 5)**

Schematics showing the placement of optic fibers in all the mice used for activating Sst$^{CeA\text{-}GPe}$ neurons with optogenetics (n = 5 mice). The CeA is colored in dark gray.
